# *Eating an elephant, one bite at a time*: predator interactions at carrion bonanzas

**DOI:** 10.1101/2023.03.31.535158

**Authors:** Aaron W. Morris, Ian Smith, Stotra Chakrabarti, Fredrick Lala, Stephen Nyagah, Joseph K. Bump

**Affiliations:** University of Minnesota, Twin Cities, MN, USA; Department of Biology, Macalester College; Wildlife Research Training Institute, Kenya

**Keywords:** camera traps, carrion ecology, intraguild interactions, interference competition, optimal foraging, predation ecology

## Abstract

Resource specific competition between predators has typically been studied from their interactions at meso-herbivore carcasses, because such carcasses are abundant. Mega-carcasses like those of elephants are rare but unparalleled in the extent of carrion biomass they offer and the long durations they can persist. These rare resource bonanzas can thus provide unique opportunities to understand sympatric species interactions within likely relaxed competitive scenarios. Using remote cameras that were operational 24-h a day, we monitored two elephant carcasses in Tsavo, Kenya, from when they were discovered until they were completely consumed or became inaccessible. While we found high temporal overlaps in activity patterns between all predators, the terrestrial predator guild (lion/leopard/spotted hyena) was not observed to feed simultaneously, suggesting strong interference competition. Based on photo-analysis and video-evidence of exclusion from a carcass, interference competition within the terrestrial predator guild favored lions over hyenas, and hyenas over leopards. The carcass at the terrestrial-aquatic interface showed more simultaneous feeding bouts between predators (crocodile/spotted hyena), indicating either facilitation and/or higher coexistence between predators that typically occupy different niches. We also observed a hippopotamus scavenging from an elephant carcass, thereby documenting a rare instance of a megaherbivore feeding on a megaherbivore. Our results highlight the importance of monitoring such carcasses through remote cameras, which can significantly add to our existing understanding of food webs and carrion ecology.

## Introduction

The carcasses of megaherbivores like elephants are nonpareil as a single source of carrion availability in their respective ecosystems, similar to *whale-falls* at ocean depths (Smith and Baco 2003). The sheer size of elephants makes them persist longer on the landscape and allows for more species to use the available carrion (Moleon et al. 2015). Apart from periods of severe droughts, elephants are not typically available as a carrion resource due to their long life and big size that helps them to avoid predation. Instead, meso-herbivore carcasses are more commonly available carrion items but only persist for significantly shorter durations before they are fully consumed (Blumenschine 1989). Consequently, most of our understanding of intraguild foraging interactions between predators/scavengers originates from interactions at these meso-herbivore carcasses and observations from megaherbivore carcasses are few.

Round the clock monitoring of megacarcasses through remote cameras provides unique opportunities to understand interactions between typically competing predators that use the carrion within (expected) relaxed competitive scenarios (when food is plenty). However, whether such resource bonanzas facilitate and/or relax intraguild interactions is an untested question. In this study, we examine the interactions within a terrestrial predator guild (between lions, spotted hyenas, and leopards) at an elephant carcass (hereafter Voi carcass), and we compare them with the interactions between a terrestrial-aquatic interface when an elephant carcass (hereafter Galana carcass) was shared between spotted hyenas and crocodiles in an East African savannah system.

## Methods

### Study Area

We conducted this research in Tsavo East National Park (TENP) within the Tsavo Conservation Area (TCA) (3°21’45.5837”, 038°35’45.9666”) in Kenya. Throughout the year, daily temperatures average between 20°C and 30°C, and this semi-arid region has two rainy seasons (Ngene et al. 2017). The region also experiences irregular severe droughts (Corfield 1973, Coe 1978). The Galana river is the only permanent river in the region, with the Tiva and Voi rivers as primary seasonal rivers (Ngene et al. 2017). In the TCA, vegetation is predominantly lowland *Acacia-Commiphora* savannah (Maingi et al. 2012), which varies in spatial and temporal densities (Gillson 2004). Wildlife diversity within the TCA includes typical savannah species such as the elephant (*Loxodonta Africana*), plains zebra (*Equus quagga*), hippopotamus (*Hippopotamus amphibius*), Nile crocodile (*Crocodylus niloticus*), lion (*Panthera leo*), spotted hyena (*Crocuta crocuta*), leopard (*Panthera pardus*), and striped hyena (*Hyaena hyaena*).

The TCA is home to Kenya’s largest population of savannah elephants (Waweru et al. 2021). The most common causes of elephant mortality in the TCA are drought (Wato et al. 2016) and poaching (Maingi et al. 2012). Due to the threat of poaching, patrol units cover the TCA intensively via ground and air. Such intensive monitoring provides for the detection of elephant carcasses quite readily.

### Field Sampling

We monitored two elephant carcasses with motion-triggered camera traps in 2019. Each carcass was monitored for the entire duration since discovery until it was completely consumed or became inaccessible. A camera trap (Cuddeback Silver) was deployed ∼15 m from each elephant carcass at a height of ∼50 cm off the ground. Cameras were programmed to take two rapid fire photographs with every trigger and set with a 5-minute delay between triggers, operational 24-h a day. Cameras were checked once a week to replace memory cards and batteries and also reposition them if necessary. We also set one Reconyx Hyperfire camera at each carcass to record videos of ensuing interactions between scavengers/predators.

### Photo-Analysis

All photographs were date and time stamped. For every trigger/event, we used one photograph to maintain a single-entry point in time. We generally used the first image that was captured unless the carcass was obstructed from view, when we then used the other image. For every image, we identified the species present and the total number of each species in the camera view. Images that were blurred and/or included unidentifiable species were discarded. For every image, we categorized the behavior of all visible species into five classes: *resting, standing, feeding, moving*, and *socializing*. For analysis, we used only still images from the Cuddeback cameras, although we report interaction videos from the Reconyx cameras to support our results.

After segregating by events, we analyzed temporal activity of spotted hyenas, crocodiles, lions, and leopards at the carcasses by developing probability distributions using a nonparametric kernel density estimation (Ridout and Linkie 2009). Spotted hyenas were the only species that overlapped between the two carcasses and for species-specific activity patterns, we used cumulative data points for hyenas across both carcasses. We further developed temporal activity overlap between the predators using images that only exhibited feeding behaviors to understand mutual resource use (Wang et al. 2015). To investigate spatio-temporal overlap, we further analyzed the proportion of photographs where two competing species were found to feed from the same carcass simultaneously. Based on the nature of interference competition between terrestrial predators that occupy similar niches, we expected high overall activity overlaps but low simultaneous feeding bouts.

## Results

The Voi carcass persisted from Aug 13, 2019 - Sept 17, 2019, while the Galana carcass had predator/scavenger activity between Sept 17, 2019 - Oct 4, 2019. We captured photographs across 2524 events at the two carcasses. The majority of these events included spotted hyenas (n=1721), followed by crocodiles (n=566), lions (n=215), and leopards (n=8). Among these events, we observed hyenas feeding in 1117 events, crocodiles in 220 events, lions in 192 events and leopards in two events. All species typically showed nocturnal activities (Figure 1), with a high degree of temporal overlap between them (Figure 2). Crocodiles and spotted hyenas showed an overlap of Δ□4= 0.74, similar to that of lions and spotted hyenas (Δ□4= 0.74), while leopards and hyenas (Δ□4= 0.28), along with leopards and lions (Δ□4= 0.24), overlapped much less in their temporal activities. Although these predators had considerable to moderate levels of temporal overlap between their overall feeding times, hyenas and crocodiles were seen to simultaneously share a carcass at 18% of the feeding events, leopards and hyenas at 0.13%, and we found no instances of simultaneous feeding between lions and hyenas.

**Figure 1.**
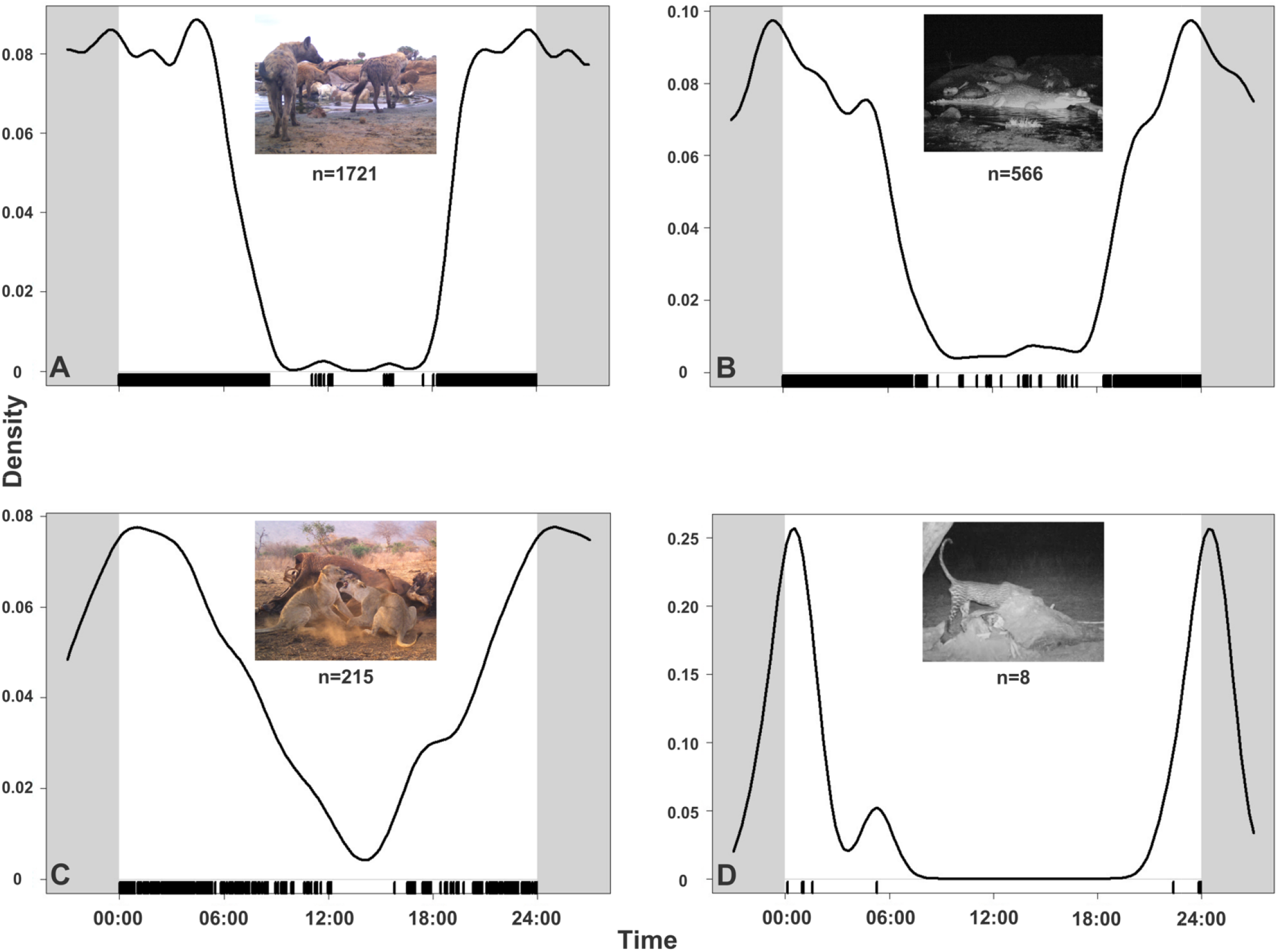
Temporal activity plots for A. spotted hyena, B. crocodile, C. lion, and D. leopard at elephant carcasses monitored through camera traps in Tsavo, Kenya. Each plot represents a kernel density of respective species appearing at the carcasses across a 24-h cycle for the entire duration a carcass was monitored. Since spotted hyenas appeared on both carcasses, we used cumulative data for them.

**Figure 2.**
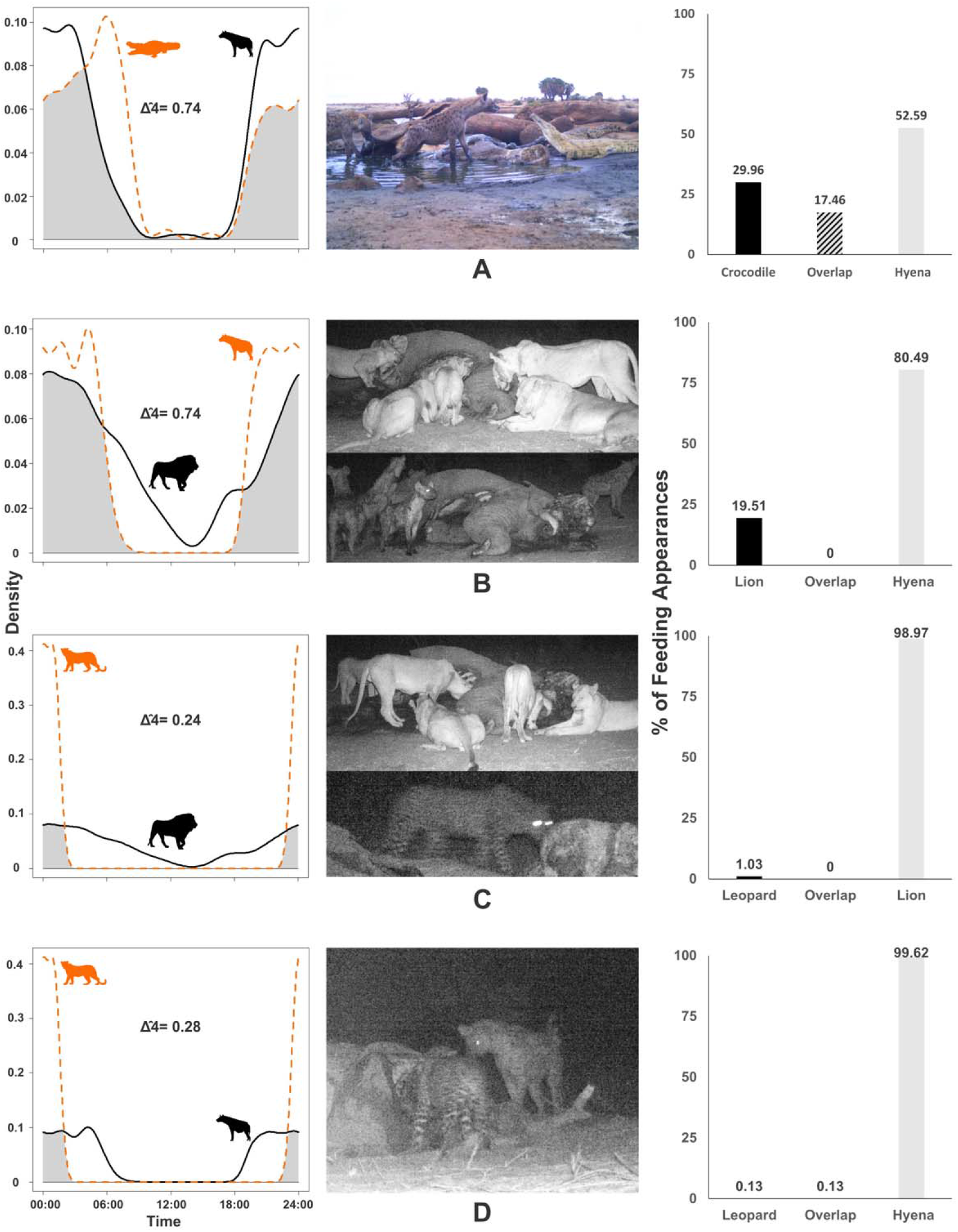
Temporal overlap between four sympatric predators in Tsavo, Kenya, feeding on elephant carcasses. A. spotted hyena:crocodile time density overlap based on events of feeding from the same carcass, image from camera trap, and percentage of photographs when the two species were recorded feeding on the carcass on their own versus together, B. spotted hyena:lion time density overlap based on events of feeding from the same carcass, image from camera trap, and proportion of photographs when the two species were recorded feeding on the carcass on their own versus together, C. lion:leopard time density overlap based on events of feeding from the same carcass, image from camera trap, and proportion of photographs when the two species were recorded feeding on the carcass on their own versus together, and D. spotted hyena:leopard time density overlap based on events of feeding from the same carcass, image from camera trap, and proportion of photographs when the two species were recorded feeding on the carcass on their own versus together.

## Discussion

Our results show high temporal overlap between competing terrestrial predators, as expected from their foraging niches and behavior. Although they were active during the same time (Figure 1), we found no instances of lions and spotted hyenas feeding from the same carcass simultaneously, and rarely did leopards and spotted hyenas do so (Figure 2). This suggests high interference competition between these three predators, although the point resource was abundant. Spotted hyenas have a broad dietary niche (able to consume meat and bones) and are in relatively high abundance in Tsavo, which may explain their first to appear and the most persistent use of the carcasses. The frequency of spotted hyenas at the Voi carcass declining when lions were present (Video 1) suggests that interference competition favors lions. Lions were also found to competitively exclude hyenas from the carcass (Video 2). Our results reflect similar interactions between this predator guild in South Africa (Amoroś et al. 2020). A leopard, although rare at the monitored carcasses, was found at a carcass simultaneously with a single spotted hyena, perhaps indicating lower levels of competition between these two species, which are of similar unit body weights. However, rivalry quickly favored the social predator among the two; a leopard was never found to be feeding when a group of hyenas was present. At the Galana carcass, crocodiles and hyenas seem to be feeding simultaneously on significantly more occasions (Figure 2). This prompts interesting questions regarding facilitation between the two predators: Do crocodiles that may not be able to open up the tough hide of an elephant carcass benefit from the presence of spotted hyenas? Or do spotted hyenas and crocodiles cope with competition at a super-abundant resource more easily because their niches are typically different? We also found an instance of a hippopotamus scavenging from the Galana elephant carcass, recording the scavenging carnivory of a megaherbivore on another megaherbivore carcass (Video 3). These instances support the benefits of monitoring megaherbivore carcasses through camera traps that record interesting and rare behaviors. African elephant carcasses may essentially act as the terrestrial analog of *whale-falls*, and further camera trap-supported research is needed across different niches to investigate one component of this possibility – how interference competition modulates carcass monopoly.

## Data Availability Statement

Since the data contains sensitive information about threatened and endangered species as well as elephant mortality, we have not made the data publicly available. However, data requests can be directed to the corresponding author directly, and we will make the data available upon request

## Acknowledgements

We would like to thank Dr. Patrick Omondi, Director, Wildlife Research & Training Institute (WRTI), Kenya for granting permissions to conduct the research. We are incredibly grateful to all the staff from WRTI from Tsavo East, and Kenya Wildlife Service rangers for assisting with fieldwork and data collection. We thank Anna Von Duyke, Elsa Heath, and Irina Kornberg for their help with data curation. SC would like to specifically thank Eleanor Michaud for inspiring the paper title.

## Author Contributions

AWM, FL & JKB conceptualized the study; AWM, FL and SN collected the data; AWM and IS curated the data; SC and IS analyzed the data; SC & AWM led the writing of the manuscript, JKB appropriated funding for the study. All authors commented and revised the drafts, and approved the final version.

